# Brain size predicts learning abilities in bees

**DOI:** 10.1101/2020.01.27.921114

**Authors:** Miguel Á. Collado, Cristina M. Montaner, Francisco P. Molina, Daniel Sol, Ignasi Bartomeus

## Abstract

A large brain is widely considered a distinctive feature of intelligence, a notion that mostly derives from studies in mammals. However, studies in insects demonstrates that cognitively sophisticated processes, such as social learning and tool use, are still possible with very small brains. Even after accounting for the allometric effect of body size, substantial variation in brain size still remains unexplained. A plausible advantage of a disproportionately larger brain might be an enhanced ability to learn new behaviors to cope with novel or complex challenges. While this hypothesis has received ample support from studies in birds and mammals, similar evidence is not available for small-brained animals like insects. Our objective is to compare the learning abilities of different bee species with their brain size investment. We conducted an experiment in which field-collected individuals had to associate an unconditioned stimulus (sucrose), with a conditioned stimulus (colored strip). We show that the probability of learning the reward-colour association was related to both absolute and relative brain size. This study shows that other bee species aside from the long studied honeybees and bumblebees, can be used in cognitive experiments and opens the door to explore the importance of relative brain sizes in cognitive tasks for insects and its consequences for species survival in a changing world.

## INTRODUCTION

A large brain is widely considered a distinctive feature of intelligence, a notion that mostly derives from studies in mammals. However, studies in insects demonstrates that cognitively sophisticated processes, such as social learning, are still possible with very small brains (Healy & Rowe, 2007). In fact, the million-fold increase in a large mammal’s brain compared to an insect brain allows mammals to have behavioral repertoires that are only just two to three times as big (Chittka & Niven, 2009). This is hardly the kind of difference expected to find between insects and mammals.

If a large behavioural repertoire is possible with a miniature brain, what benefits obtain animals by investing in larger brains? Because brain size scales allometrically with body size (Burger et al., 2019), an explanation is that biophysical constraints forces larger animals to have more and/or larger neurons (Chittka & Niven, 2009). It is for instance easy to imagine that the bigger muscles of larger animals will require greater numbers of motor neurons and axons with larger diameters to cover longer distances (Chittka & Niven, 2009). More neurons may also allow greater replication of neuronal circuits, adding precision to sensory processes, detail to perception, more parallel processing and enlarged storage capacity (Niven & Laughlin, 2008). These explanations are however insufficient because substantial variation in brain size remains when the allometric effect of body size is taken out (Gonda et *al.*, 2013). Given that neural tissue is extremely costly to maintain, what is the purpose of expanding the brain?

A plausible advantage of a disproportionately larger brain might be an enhanced ability to learn new behaviors to cope with novel or complex challenges (Sol et *al.*, 2008). While this hypothesis has received ample support from studies in birds and mammals, similar evidence is not available for small-brained animals like insects. Our insufficient understanding of the benefits of miniature brains remains thus a major obstacle for a general theory of brain evolution, and even cast doubts on whether variation in brain size is biologically meaningful (Healy & Rowe, 2007).

In this study, we try to address this gap with an experimental comparative analysis in bees. Bees have historically fascinated biologists because of their small nervous systems compared to the complexity and diversity of their behavior (Chittka & Niven, 2009; Vasas & Chittka, 2018). Numerous species, including bees, are reported to be able to create memories of rewarding experiences (Matsumoto & Mizunami 2000; Menzel, 1999; Daly & Smith, 2000) as well as of punishment (Vergoz et *al.*, 2007), and those memories can be retrieved at different times after learning, both in the short- and in the long-term (Giurfa, 2015). Species have also substantially diversified in brain size despite sharing the same brain architecture. For example, bigger relative brain sizes have been suggested to be related to diet specialism in bees, as specialist need to locate and remember target floral resources (Sayol et *al.*, 2020).

We measured learning abilities in wild individuals of 32 bee species captured in the wild, and then used a phylogenetic comparative framework to test whether species that performed better in the learning task had larger absolute and/or relative brains. Learning abilities were measured with a novel quick-to-perform experimental method proposed by Muth et *al.*, 2017 to assess speed at colour learning. Associative learning is a highly developed cognitive ability in bees with substantial ecological relevance. Importantly, associative learning assays are short enough to be suitable for highly stress-intolerant species and facilitate as well standardization across species with varying life histories and ecologies, two major obstacles hindering past progress in linking brain size and learning improvement. The possibility to perform experiments directly from individuals captured in the field allows experimentation with non-model taxa, providing opportunities for broader comparative analyses of cognition.

## MATERIAL AND METHODS

### Study subjects

We opportunistically captured bees by hand netting (n = 202 individuals) from March to June 2018 in different open fields and urban parks from Andalusia, South of Spain. Bees were kept individually in vials in cold storages and transported to the laboratory, where they were transferred into separated transparent plastic enclosures for the behavioral assay (Fig. 1), which was conducted within the following three hours after capture. After the assays, all individuals were identified at the species level by a taxonomist (F. P. M.), yielding a sample of 32 unique species from 14 genera (Table S2).

**Figure 1.**
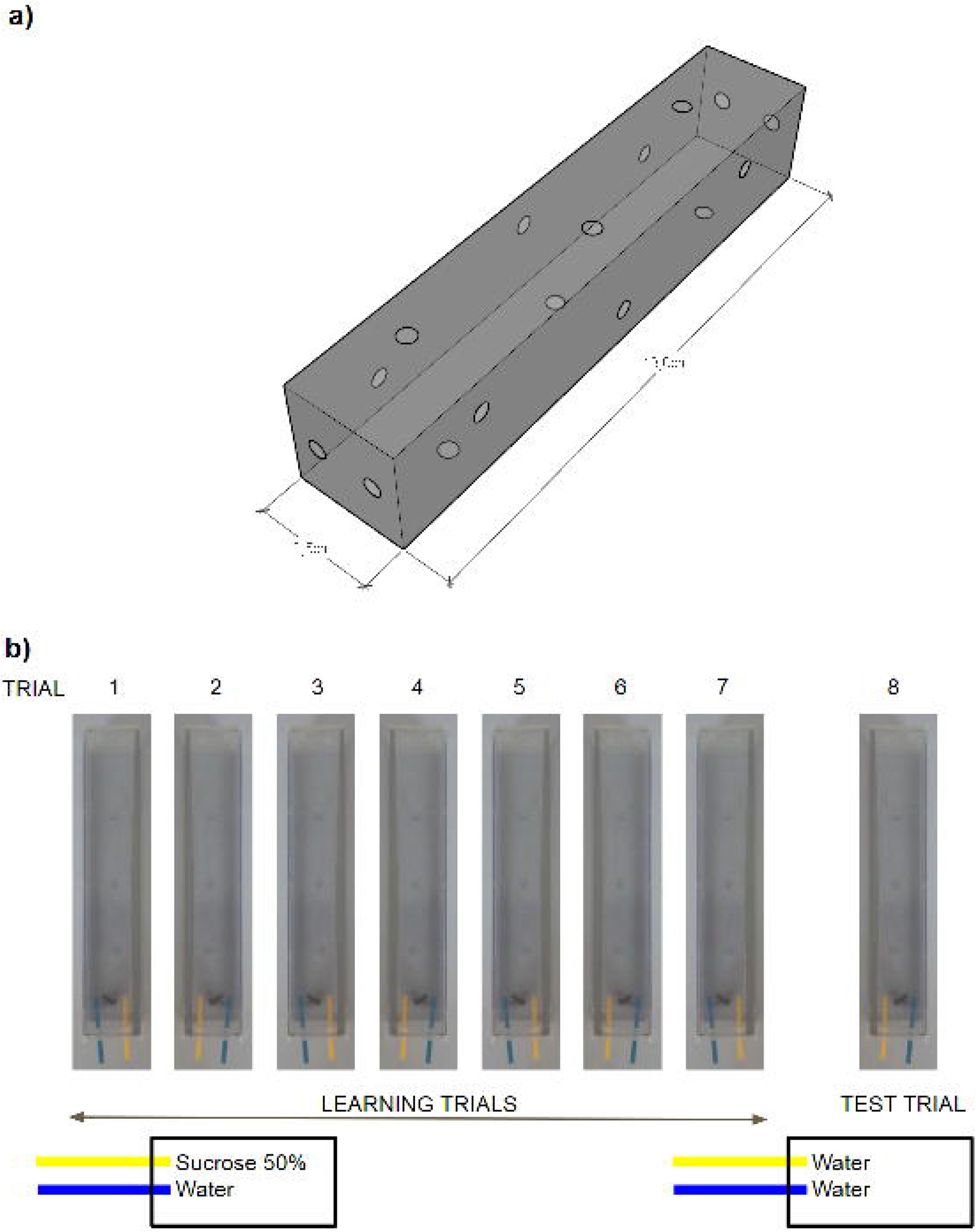
The complete experimental display. a) PVC experimental enclosures used for the experiment (2.5 cm width and 13 cm long). Multiple holes were done for air circulation and easy strip offering from both extremes. b) An example of the sequence of one complete set of assays for one single individual, where one color is associated with a reward and it is maintained until the final test trial, where both strips are unrewarded.

### Experimental assay

Before starting the learning assays, bees were left 30 minutes in the individual enclosures to allow them to awake from the cold and habituate to the experimental conditions. The experimental enclosures were built by the attachment of two 2.5 cm transparent PVC angles with ventilation holes, with removable perforated lids attached at both extremes (Fig. 1a). The experimental enclosures rested during the experiments over a grey surface to avoid external stimulus. Associative learning was measured by a multi-choice free-moving proboscis extension protocol (FMPER, modified from Muth et *al.* 2017), where the animal had to learn to associate a reward (50% sucrose) with an arbitrary stimulus (a color). Each experimental trial consisted in the presentation of a yellow and blue cardboard small strips (3 × 0.2 cm) easily distinguishable by bees vision (Chittka & Wells, 2004). The strips were presented always at the opposite extreme from where the individual was staying in the enclosure. One of the colored strips was dipped in sucrose and the other one in water. The color with the reward was randomly chosen but maintained during the whole experiment for each individual. The assay started when the individual reached the middle of the enclosure in its way towards the strips. We measured the time needed for the individual to reach the strips and extend its proboscis to start drinking on one of them, waited for 3 seconds and removed the strip (Fig. 1b). We allowed the bee to explore the remaining strip and again removed it after the individual had drunk for 3 seconds. Once this exploratory trial ended, the process was repeated every 10 minutes six more times to allow individuals to associate the stimuli (color) with the reward (food) through operant conditioning (training trials), switching the strips position to avoid confounding color with position (Fig. 1b). The only modification from the original FMPER protocol (Muth et *al.*, 2017) is that we removed the first acclimatization trials and considered the first trial as acclimatization/exploration.

A trial was considered successful when the bee chose first the strip with sucrose and unsuccessful otherwise. The trial was considered finished when the subject drank from both strips or otherwise capped after two minutes. After seven training trials, we tested whether the individual had learned to associate a color with a reward by means of a unrewarded test, where both strips were dipped into water. learning improvement was defined in terms of success or failure in solving the test. When solved, we also quantified the time needed to start drinking from the right strip.

In general, and taking into account the high stressant experimental environment, bees responded well to the experimental procedure, especially those from *Andrena sp.*, *Apis mellifera*, *Bombus sp.*, *Lasioglossum sp.*, and *Rhodanthidium sticticum*. However, forty-five individuals from *Anthophora sp.*, *Eucera sp.* and *Xylocopa sp.* either ignored or did not react to a full experimental procedure (Table S2). Consequently, these species were not used in the analyses.

### Brain measurements

After the experiment, bees were anesthetized in cold chambers (Crook, 2013) and decapitated. The head was fixed in 4% paraformaldehyde with phosphate buffer saline (PBS). The fixative solution embedded the brain and dehydrated the tissue, preventing brain from degradation for a long period. Brains were extracted from the head capsule, separated from the tracheas and fat bodies to avoid weighting errors, and placed on a small piece of tared Parafilm®. Fixative solution was dried from the brain using Kimwipes® tissues and then the brain was weighted in a microbalance to microgram accuracy (Sartorius Cubis®). Brain weight was used as a proxy of brain size, as it is strongly correlated with brain volume of the mushroom bodies (correlation coefficient = 0.85; p-value < 0.001, Sayol et *al.*, 2020), which are the neuropil centers of most cognitive abilities (Dujardin, 1850). Body size was measured as the inter-tegular distance, that is, the distance between the wing bases, usually used as a proxy of body size in bees (Kendall et *al.*, 2019).

### Data analysis

We used Bayesian phylogenetic generalised linear mixed models (PGLMM), as implemented in the package *brms* (Bürkner, 2017). Specifically, we analysed whether the probability of success during the training trials (i.e. the correct strip is chosen) increased over time and whether for those trials that were successful the time to start feeding (i.e. latency) declined over time. To model success / failure we used models with Bernoulli error structures, while to model latencies we used Gaussian error structures. In all models, species were treated as a random factor and we incorporate a phylogenetic covariance matrix to control for the influence of phylogeny. The phylogeny used was a maximum-likelihood phylogenetic tree of the superfamily Apoidea at the genera level (Hedtke et *al.*, 2013). Due to the absence of infrageneric phylogenies for our genera, we simulated infrageneric polytomies within our phylogeny. Species tips were added to the phylogenetic tree genera nodes as polytomies of equal branch length relative to the genera branch length (Kendall et *al.*, 2019) using the *phytools* package (version 0.6-44; Revell, 2012). Intra-class coefficients (ICC) were used to validate the assumption of comparative approach that variation in learning ability across species is higher than within species. ICC is a descriptive statistic, ranging from zero to one, that can be used when quantitative measurements are made on units that are organized into groups, in this case different measurements within species grouped by phylogenetic relationships. A value near zero is interpreted as the variable is not explained by the phylogenetic relationships, and a value near one means that the variable is better explained by intrinsic characteristics of the phylogeny. Finally, we also examined using PGLMM whether individuals that better learned to associate color and reward during the training period exhibited a higher probability of successfully solving the unrewarded stimulus test than those that did not learn it.

To test the prediction that large brains enhance learning abilities, we modelled the success in the test and the learning improvement as a function of brain size using the PGLMM framework described above. Learning improvement was defined as one minus the division between the latency to touch the correct cue the first time and the latency to touch the correct cue in the test (e.g. values near one mean larger learning improvements). Following previous studies, we analyzed brain size both in absolute terms (brain weight) and relative to body mass, fitting separate models for each brain size metric: absolute brain size (log-transformed) and relative brain size estimated as brain size residuals (Wurm & Fisicaro, 2014). These residuals were extracted from a log-log regression of brain size against body size (LM estimate ± SE = 2.07 ± 0.08, p < 0.001, R^2^= 0.82). To avoid that innate preferences systematically bias the experiment, we randomized position and colour associations with the reward.

## RESULTS

Most bees learned to associate a color with a food reward. More than half of the individuals that reached the test drank from the correct strip (66%). This was higher than expected by chance (i.e. 50%; Chi-squared goodness of fit 10.8, p = 0.001). Time to touch the correct strip decreased along the trials (PGLMM Negative Binomial, β = −0.13 ± 0.02, IC = −0.16 – −0.10, ICC: 0.28, Fig. 2, Table 1) and bees had more chances of success in the later trials than in the earlier ones (PGLMM Negative Binomial β = 0.07 ± 0.02, IC = 0.03 − 0.10, ICC: 0.03). Removing the first trial, which can be merely exploratory, did not change the results. Finally, those with higher number of learning successes during the whole experimental process were more likely to pass the learning test (PGLMM Bernoulli β = 0.74 ± 0.16, IC = 0.45 – 1.06, ICC = 0.17, Table 1). The probability of learning success and learning improvement were poorly explained by phylogeny as shown by the low ICC values reported above. Learning improvement had a slightly higher phylogenetic signal (ICC: 0.38-0.39), maybe indicating that certain groups, can be better at learning.

**Table 1.** Results of the Bayesian models of learning, and learning related to brain size. IC = Interval Confidence, ICC = Intra-class coefficient, β = Estimate ± Standard Error.

| Formula | $\beta$ | IC | ICC (Species) | Notes |
| --- | --- | --- | --- | --- |
| **LEARNING** |  |  |  |  |
| Time until touching the correct strip ~ Trial number | $-0.13 \pm 0.02$ | $-0.16 - -0.10$ | 0.28 | All trials considered |
| Trial success ~ Trial number | $0.07 \pm 0.02$ | $0.03 - 0.10$ | 0.03 | All trials considered |
| Number of trial success ~ Trial number | $0.74 \pm 0.16$ | $0.45 - 1.06$ | 0.17 | First trial removed |
| **LEARNING COMPARED WITH BRAIN SIZES** |  |  |  |  |
| Learning success ~ Absolute brain size | $0.79 \pm 0.27$ | $0.29 - 1.38$ | 0.10 | Just learning test |
| Learning success ~ Relative brain size | $1.26 \pm 0.78$ | $-0.26 - 2.80$ | 0.24 | Just learning test |
| Learning improvement ~ Absolute brain size | $0.05 \pm 0.08$ | $-0.13 - 0.21$ | 0.39 | Just learning test |
| Learning improvement ~ Relative brain size | $0.21 \pm 0.27$ | $-0.32 - 0.75$ | 0.38 | Just learning test |

**Figure 2.**
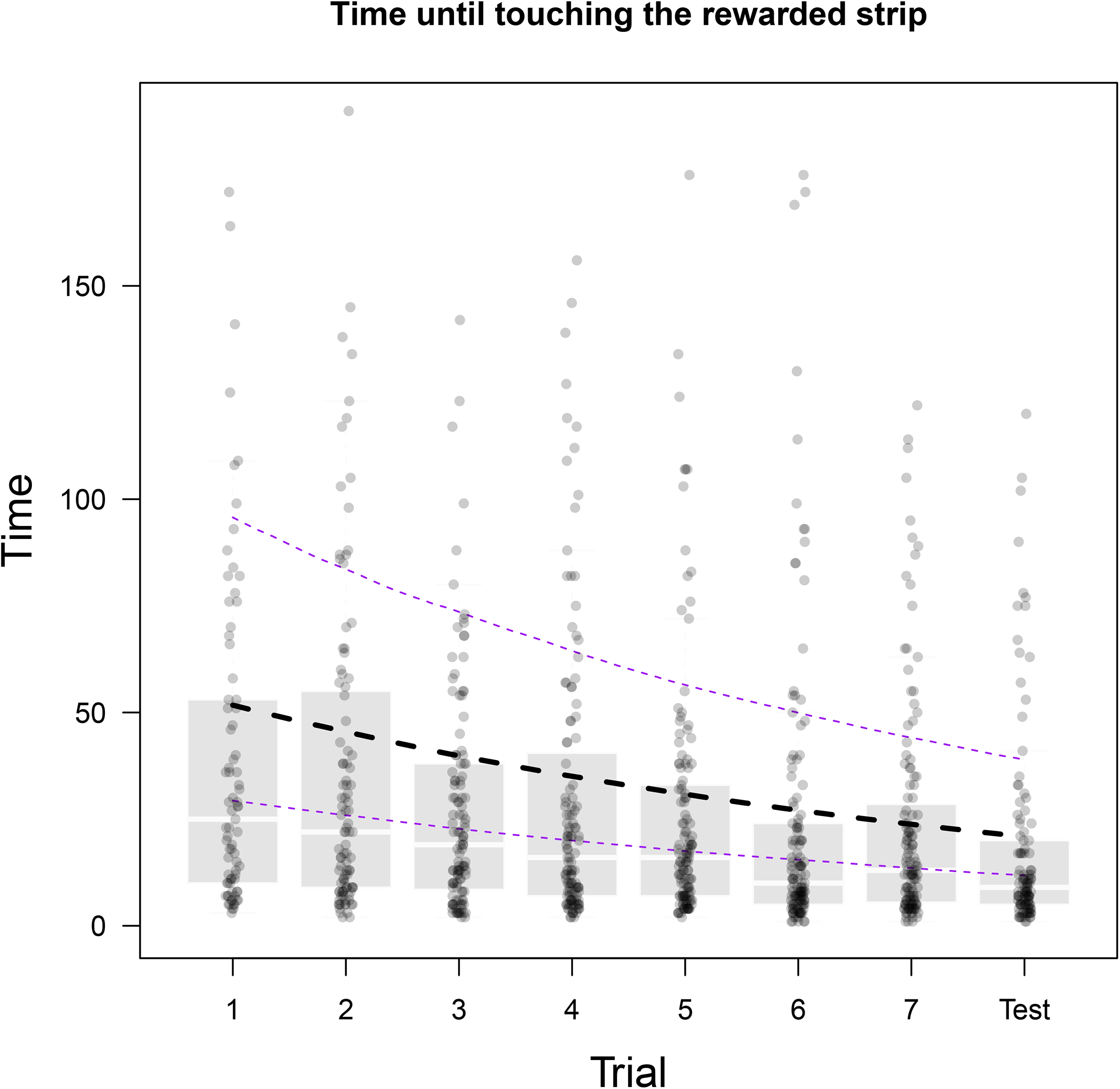
Time (seconds) until touching the rewarded strip decreased along trials. Dots represent each individual success. Boxplots represent from Q2 to Q3 of those success for each trial, with the median drawn in white. Overlapping the boxplot is the estimate and confidence intervals of the PGLMM negbinomial model (β = −0.13 ± 0.02, IC = −0.16 – −0.10, ICC: 0.28). Last trial was considered the test as it was not rewarded.

**Figure 3.**
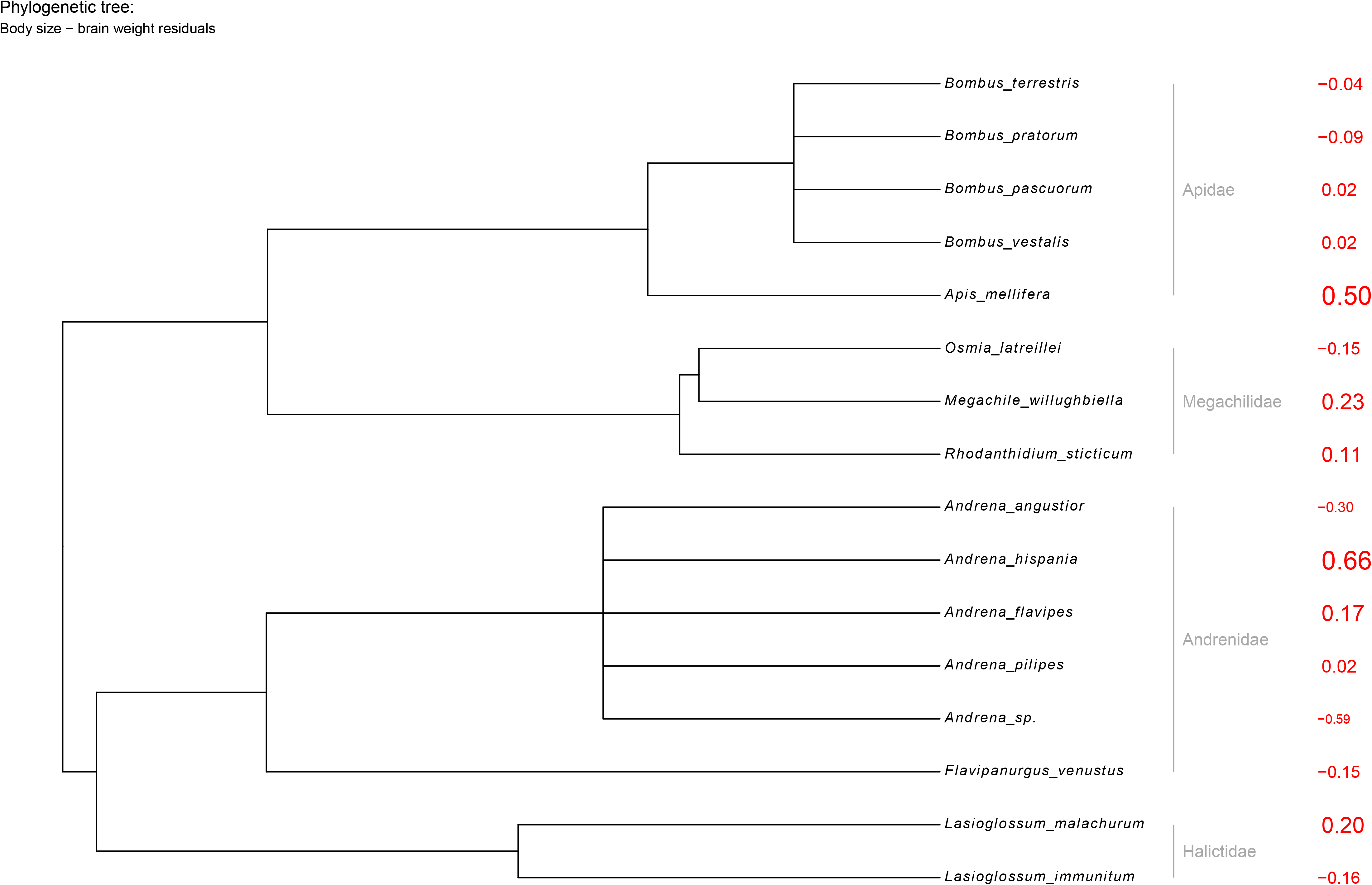
Phylogenetic tree of the studied species used for the Bayesian analysis to control whether models were explained by the phylogenetic group heritage. Numbers in red represent the body size - brain size residuals for each species. Values above zero have bigger brains than expected by their body sizes, and values below zero have smaller brain sizes.

Bees with bigger brains were more likely to succeed in the learning test. Success in the learning test was positively correlated with both absolute brain size (PGLMM Bernoulli β = 0.79 ± 0.27, IC = 0.29 – 1.38, ICC: 0.10, Fig. 4a) and with brain residuals (PGLMM Bernoulli β = 1.26 ± 0.78, IC = −0.26 – 2.80, ICC: 0.24, Figure 4b). In addition, for the subset of individuals that succeed in the learning test, individuals that experienced a larger learning improvement along trials had slightly bigger brains, both in absolute terms (PGLMM Bernoulli β = 0.05 ± 0.08, IC: −0.13 – 0.21, ICC: 0.39, Figure 4c, Table 1) and relative to body size (PGLMM Bernoulli, β = 0.21 ± 0.27, IC: −0.32 – 0.75, ICC: 0.38, Figure 4d, Table 1), but this effect is highly variable.

**Figure 4.**
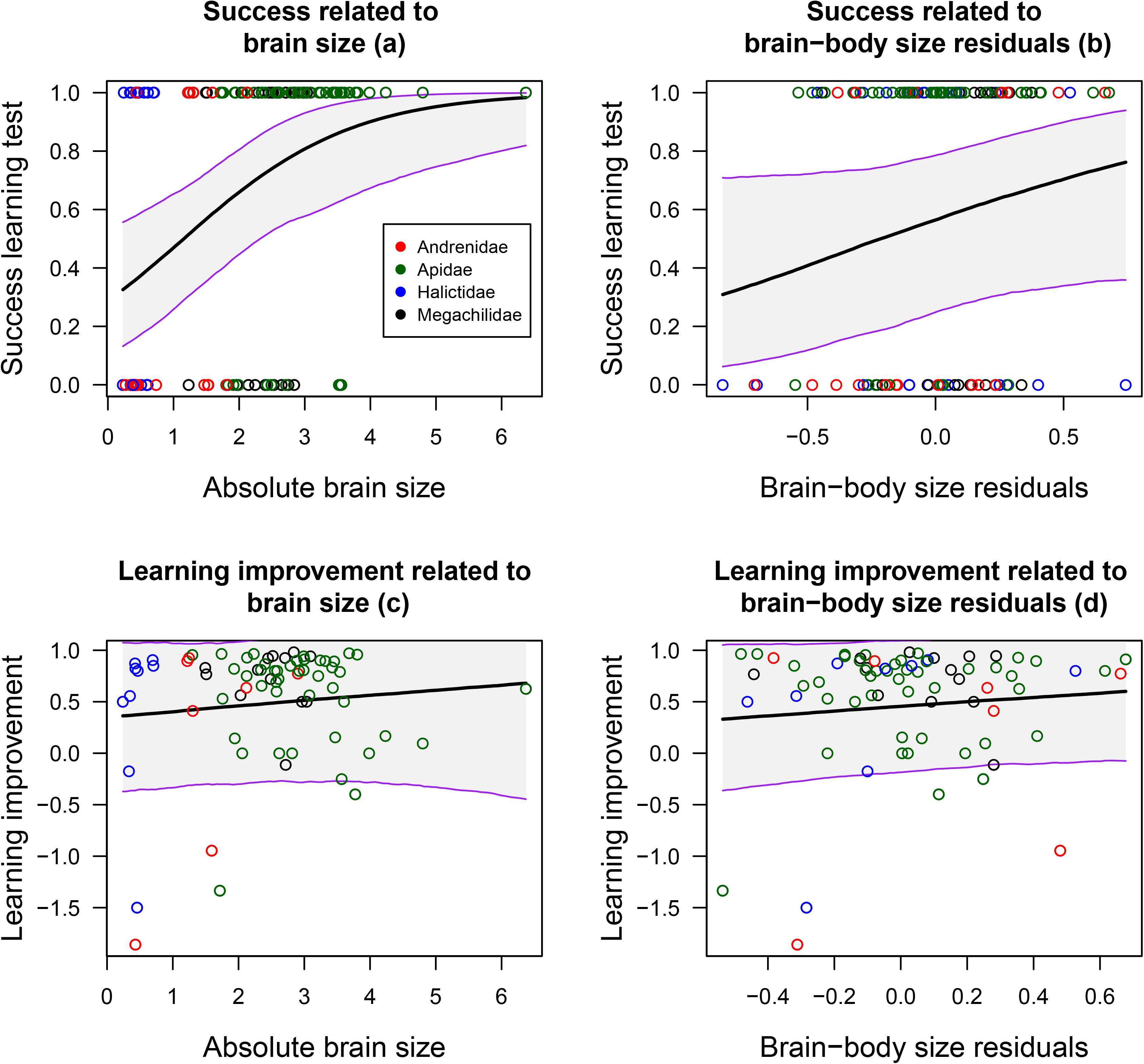
Correlation between brain size and learning improvement. a) Increasing probability (from zero to one) of success with bigger brain sizes (brain weight in mg). b) Increasing probability of success with bigger relative brain sizes. c) The correlation between the learning improvement from the first trial to the last test with absolute brain sizes. d) The correlation between learning improvement and relative brain sizes. Overlapping lines represent the trends extracted from the PGLMM models and their IC.

## DISCUSSION

Highly controlled laboratory experiments allow testing detailed learning abilities, but only for a handful of species (e.g. honeybees, bumblebees, cockroaches, fruit flies) that can be raised in laboratory conditions. For example, some of these experiments often involve stressful conditions, like individuals being fully harnessed in Proboscis Extension Reflex (PER) protocols (Takeda, 1961). Therefore, laboratory experiments are only suitable for highly stress-tolerant species. We chose a modified version of the multi-choice free-moving proboscis extension protocol (FMPER, modified from Muth et *al.*, 2017) to test simple learning abilities simultaneously for several bee species never tested before. This novel quick-to-perform experimental method allowed us to conduct a comparative analysis including multiple bee species captured directly from the field. We found that most bee species, including solitary species never used before in cognitive experiments, can learn to associate a color with a reward. Interestingly, species differed in their learning abilities, and these differences were in part explained by brain size. Thus, the probability to learn increased with both absolute and relative brain sizes, while learning improvement along the trials was weakly related to brain size, but was also influenced by the species position in the phylogeny.

Expanding the range of species evaluated by applying quick-to-perform experimental protocols to wild species can provide important insight into the relationship between brain size and learning. Thus, although our experiment was not designed to analyze differences across social and solitary species, we observed that some solitary species can show similar learning abilities than social species. This appears to contradict the traditional view that social bees have more complex learning abilities than solitary bees (Reader & Laland, 2002), although more research is needed to properly confirm it.

We used in our analysis both relative brain size and absolute brain size, two metrics with different assumptions and interpretations (Burger et al., 2019; Healy & Rowe, 2007). Absolute brain size was clearly related to learning abilities, this implies that just having a larger amount of neural tissue can improve learning abilities, at least within our studied taxa. However, larger bees also have larger visual organs and mobility, which can facilitate the learning task for non-cognitive reasons. Interestingly, having a bigger brain than expected by their body size was also correlated with learning abilities. This suggests that investing in brain size payoff in terms of learning abilities (but see Healy & Rowe, 2007; Chittka & Niven, 2009). Note that the ICC values indicate that phylogeny is important in explaining some of these relationships, as body size is strongly constrained phylogenetically. In that way, all bumblebee species considered are the ones with larger absolute brain sizes, but when relative brain size is considered, they are well distributed across the data. Our measure of learning improvement was specially phylogenetically constrained, therefore we suspect that other conserved traits linked to body size, such as mobility may better explain this variable.

A handful of species did not react to the experimental settings, showing no interest for the colored stripes. Specifically, we found 11 species that did not react to any complete experimental protocol (i.e. were not active in enough trials to consider a valid test, etc.) and six more species that fully ignored the experimental setting. Despite the experiment was designed to isolate learning, there are other confounding variables that can affect the experimental responses including: stress management (Even et *al.*, 2012), neophobia (Forrest & Thomson, 2008), motivation (Dyer et *al.*, 2002) or color perception (Chittka & Wells, 2004). Therefore, species that did not react were not necessarily “species unable to learn” and alternative explanations are possible. Using wild animals can also have caveats, as stress may make individuals to behave in strange ways. However, evidence from *Apis mellifera* does not suggest that using wild individuals change the results of learning tests compared to bees born in laboratory (Muth et *al.*, 2017), but further analyses are needed for species that do not habituate so well to experimental settings. Despite not all species are suitable for such experimental approach, the number of candidate species we can test increases dramatically in comparison to classic experimental settings such as the Proboscis Extension Reflex mostly done in *A. mellifera* and bumblebees (Takeda, 1961; Vergoz et al., 2007; Laloi et al., 1999)

Our results provide the first evidence that insects with larger brains, both in absolute and relative terms, perform better in an associative learning test than species with smaller brains, challenging previous claims that variation in brain size is not biologically meaningful (Chitka & Niven, 2009; Healy & Rowe 2007). However, it remains to be demonstrated whether similar patterns can be extended to other learning mechanisms. The underlying processes also warrant explanation. The challenge is to elucidate whether variation in learning speed across species reflects sensorial, cognitive, physical or emotional responses, and how these responses are associated with finer brain structures like mushroom bodies, neuron density or optimized neurons synapses.

## AKNLOWLEDGEMENTS

We wanted to thank Felicity Muth & Anne S. Leonard for their inspiring work on FMPER. Carlos Zaragoza, for his help doing some of the experiments. We are grateful to Consejería de Medio Ambiente, Junta de Andalucía, for permission to work in Sierra de Cazorla and providing invaluable facilities there, and to EBD-CSIC for making Roblehondo field station available to us.We are also grateful to MINISTERIO DE ECONOMÍA Y COMPETITIVIDAD, GOBIERNO DE ESPAÑA for the projects Grant/Award Numbers: CGL2013-47448-P and CGL2017-90033-P.

