## Supplementary material for "Brain size predicts learning abilities in bees": Table S1

**Table S1** Names and coordinates from the sample sites where the individuals were captured.

| Site | Latitude | Longitude |
|---|---|---|
| Alamillo Park | 37,42 | -5,99 |
| El Arboreto, botanical garden | 37,39 | -6,04 |
| Pablo de Olavide campus | 37,35 | -5,94 |
| Aznalcázar, natural park | 37,23 | -6,17 |
| La Rocina, Doñana natural park | 37,12 | -6,51 |
| Niebla, plot | 37,41 | -6,68 |
| Convento de la luz, plot | 37,29 | -6,75 |
