## Supplementary material for "Brain size predicts learning abilities in bees": Table S2

The list of the species used for the experiment, with the minimum time to respond to the first trial (the trial considered exploratory), the proportion of succeeders in the test compared to the number of individuals that made it to the test, and the number of captured individuals that started the test. Unidentified species where assigned to unique morphospecies.

| Species | Minimum reaction time to the first trial (seconds) | Proportion of individuals that succeeded in the test | Captured individuals |
|---|---|---|---|
| *Bombus pratorum* | 3 | 2/4 | 5 |
| *Apis mellifera* | 4 | 4/4 | 7 |
| *Bombus terrestris* | 4 | 35/41 | 43 |
| *Andrena morpho1* | 6 | 0/2 | 15 |
| *Bombus pascuorum* | 7 | 4/5 | 5 |
| *Lasioglossum malachurum* | 7 | 4/8 | 12 |
| *Rhodanthidium sticticum* | 7 | 13/21 | 26 |
| *Andrena angustior* | 10 | 1/3 | 3 |
| *Andrena pilipes* | 11 | 3/6 | 7 |
| *Andrena flavipes* | 18 | 2/4 | 6 |
| *Andrena hispania* | 22 | 1/1 | 1 |
| *Lasioglossum immunitum* | 29 | 7/15 | 18 |
| *Bombus (Psithyrus) vestalis* | 53 | 1/1 | 1 |
| *Flavipanurgus venustus* | 31 | 0/2 | 5 |
| *Osmia latreillei* | NA | 1/2 | 5 |
| *Megachile willughbiella* | NA | 0/0 | 1 |
| **Species that did not respond to any complete set of trials** | | | |

| | | | |
|---|---|---|---|
| *Xylocopa cantabrita* | 47 | 0/0 | 6 |
| *Andrena cinerea* | 53 | 0/0 | 2 |
| *Panurgus dargius* | 60 | 0/0 | 1 |
| *Lasioglossum morpho1.* | 138 | 0/0 | 4 |
| *Eucera morpho1.* | 141 | 0/0 | 2 |
| *Andrena rhyssonota* | NA | 0/0 | 7 |
| *Andrena labialis* | NA | 0/0 | 1 |
| *Eucera elongatula* | NA | 0/0 | 2 |
| *Osmia caerulescens* | NA | 0/0 | 2 |
| *Panurgus morpho1.* | NA | 0/0 | 1 |
| **Species that did not respond to any trial** | | | |
| *Eucera rufa* | NA | 0/0 | 1 |
| *Eucera notata* | NA | 0/0 | 4 |
| *Anthophora plumipes* | NA | 0/0 | 1 |
| *Anthophora dispar* | NA | 0/0 | 1 |
| *Xylocopa violacea* | NA | 0/0 | 4 |
| *Anthophora retusa* | NA | 0/0 | 2 |
